# Potential role of the *X* circular code in the regulation of gene expression

**DOI:** 10.1101/2020.03.23.003251

**Authors:** Julie D. Thompson, Raymond Ripp, Claudine Mayer, Olivier Poch, Christian J. Michel

## Abstract

The *X* circular code is a set of 20 trinucleotides (codons) that has been identified in the protein-coding genes of most organisms (bacteria, archaea, eukaryotes, plasmids, viruses). It has been shown previously that the *X* circular code has the important mathematical property of being an error-correcting code. Thus, motifs of the *X* circular code, i.e. a series of codons belonging to *X*, which are significantly enriched in the genes, allow identification and maintenance of the reading frame in genes. *X* motifs have also been identified in many transfer RNA (tRNA) genes and in important functional regions of the ribosomal RNA (rRNA), notably in the peptidyl transferase center and the decoding center. Here, we investigate the potential role of *X* motifs as functional elements in the regulation of gene expression. Surprisingly, the definition of a simple parameter identifies several relations between the *X* circular code and gene expression. First, we identify a correlation between the 20 codons of the *X* circular code and the optimal codons/dicodons that have been shown to influence translation efficiency. Using previously published experimental data, we then demonstrate that the presence of *X* motifs in genes can be used to predict the level of gene expression. Based on these observations, we propose the hypothesis that the *X* motifs represent a new genetic signal, contributing to the maintenance of the correct reading frame and the optimization and regulation of gene expression.

**Author Summary:** The standard genetic code is used by (quasi-) all organisms to translate information in genes into proteins. Recently, other codes have been identified in genomes that increase the versatility of gene decoding. Here, we focus on the circular codes, an important class of genome codes, that have the ability to detect and maintain the reading frame during translation. Motifs of the *X* circular code are enriched in protein-coding genes from most organisms from bacteria to eukaryotes, as well as in important molecules in the gene translation machinery, including transfer RNA (tRNA) and ribosomal RNA (rRNA). Based on these observations, it has been proposed that the *X* circular code represents an ancestor of the standard genetic code, that was used in primordial systems to simultaneously decode a smaller set of amino acids and synchronize the reading frame. Using previously published experimental data, we highlight several links between the presence of *X* motifs in genes and more efficient gene expression, supporting the hypothesis that the *X* circular code still contributes to the complex dynamics of gene regulation in extant genomes.

## Introduction

Codes are ubiquitous in genomes: for example, the genetic code [1], the nucleosome positioning code [2], the histone code [3], the splicing code [4], mRNA degradation code [5], or the ‘codon usage’ code [6], to name but a few. The standard genetic code [1] is probably the most well-known genome code, and represents one of the greatest discoveries of the 20th century. All known life on Earth uses the (quasi-)same triplet genetic code to control the translation of genes into functional proteins. The fact that there are 64 possible nucleotide triplet combinations but only 20 amino acids to encode, means that the genetic code is redundant and most amino acids are encoded by more than one codon. This redundancy allows for the encoding of supplementary information in addition to the amino acid sequence [7-9], and significant efforts have been applied recently to understand the multiple layers of information or ‘codes within the code’ [10] that can be exploited to increase the versatility of genome decoding.

Here, we focus on an important class of genome codes, called the circular codes, first introduced by Arquès and Michel [11] and reviewed in [12,13]. In coding theory, a circular code is also known as an error-correcting code or a self-synchronizing code, since no external synchronization is required for reading frame identification. In other words, circular codes have the ability to detect and maintain the correct reading frame. For example, comma-free codes are a particularly efficient subclass of circular codes, where the reading frame is detected by a single codon. The genetic code was originally proposed to be a comma-free code in order to explain how a sequence of codons could code for 20 amino acids, and at the same time how the correct reading frame could be retrieved and maintained [14]. However, it was later proved that the modern genetic code could not be a comma-free code [15], when it was discovered that *TTT*, a codon that cannot belong to a comma-free code, codes for phenylalanine. Other circular codes are less restrictive than comma-free codes, as a frameshift of 1 or 2 nucleotides in a sequence entirely consisting of codons from a circular code will not be detected immediately but after the reading of a certain number of nucleotides.

By excluding the four periodic codons {*AAA,CCC,GGG,TTT*} and by assigning each codon to a preferential frame (i.e. each codon is assigned to the frame in which it occurs most frequently), a circular code was identified in the reading frame of protein coding genes from eukaryotes and prokaryotes [11,16]. This so-called *X* circular code consists of 20 codons (Fig 1)

**Fig 1.**
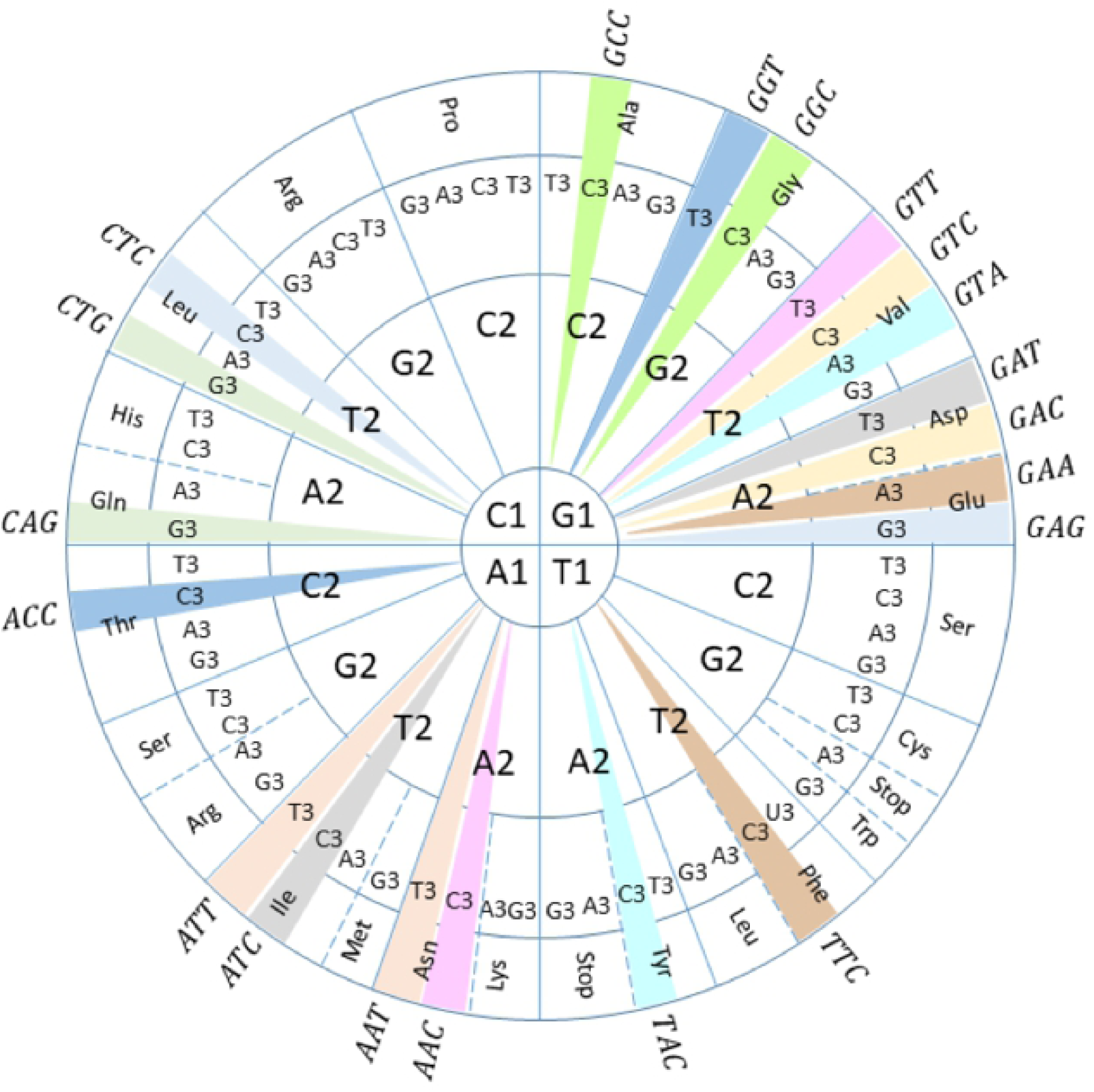
Circular representation of the genetic code, adapted from [19], with the 20 codons of the *X* circular code shown on the circumference. The numbers after the nucleotides indicate their position in the codon. *X* codons that are complementary to each other are highlighted in the same color.

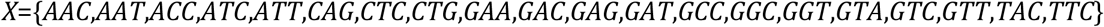

and codes for the following 12 amino acids (three and one letter notation):

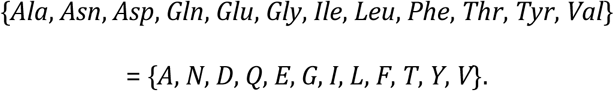

Other circular codes, and notably variations of the common *X* circular code, are hypothesized to exist in different organisms [16-18].

The *X* circular code has important mathematical properties, in particular it is self-complementary [11], meaning that if a codon belongs to *X* then its complementary trinucleotide also belongs to *X*. Like the comma-free codes, the *X* circular code also has the property of synchronizability. It has been shown that, in any sequence generated by the *X* circular code, at most 13 consecutive nucleotides are enough to always retrieve the reading frame [11]. In other words, any sequence ‘motif’ containing 4 consecutive *X* codons is sufficient to determine the correct reading frame (Fig 2) and [20]. More formal definitions of the mathematical properties (theorems) of the *X* circular code are available in a number of reviews [12-13,21].

**Fig 2.**
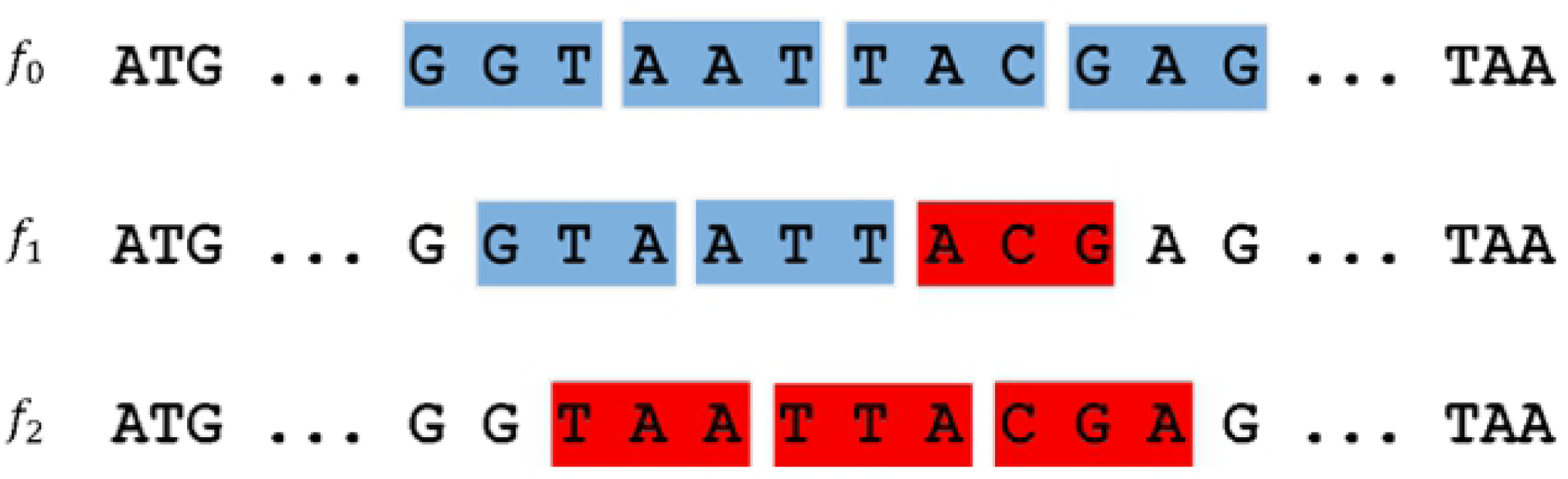
Retrieval of the reading frame in a *X* motif constructed with the *X* circular code. Codons belonging to the *X* circular code are indicated in blue, while non-*X* codons are shown in red. Among the three possible frames, only the reading frame 0 contains codons of the *X* circular code exclusively.

The hypothesis of the *X* circular code in genes is supported by evidence from several statistical analyses of modern genomes. We previously showed in a large-scale study of 138 eukaryotic genomes that *X* motifs (defined as series of at least 4 codons from the *X* circular code) are found preferentially in protein-coding genes compared to non-coding regions with a ratio of ∼8 times more *X* motifs located in genes [22]. More detailed studies of the complete gene sets of yeast and mammal genomes confirmed the strong enrichment of *X* motifs in genes and further demonstrated a statistically significant enrichment in the reading frame compared to frames 1 and 2 (*p*-value<10^−10^) [23-24]. In addition, it was shown that most of the mRNA sequences from these organisms (e.g. 98% of experimentally verified genes in *S. cerevisiae*) contain *X* motifs. Intriguingly, conserved *X* motifs have also been found in many tRNA genes [25], as well as in important functional regions of the 16S/18S ribosomal RNA (rRNA) from bacteria, archaea and eukaryotes [26-27], which suggest their involvement in universal gene translation mechanisms. More recently, a circular code periodicity 0 modulo 3 was identified in the 16S rRNA, covering the region that corresponds to the primordial proto-ribosome decoding center and containing numerous sites that interact with the tRNA and messenger RNA (mRNA) during translation [20].

Based on these observations, it has been proposed that the *X* circular code represents an ancestor of the modern genetic code that was used to code for a smaller number of amino acids and simultaneously identify and maintain the reading frame [27]. Intriguingly, the theoretical minimal RNA rings, short RNAs designed to code for all coding signals without coding redundancy among frames, are also biased for codons from the *X* circular code [28]. These RNA rings attempt to mimic primitive tRNAs and potentially reflect ancient translation machineries [29-30]. The question remains of whether the *X* motifs observed in modern genes are simply a vestige of an ancient code that might have existed in the early stages of cellular life, or whether they still play a role in the complex translation systems of extant organisms.

In this work, we define a (very) simple density parameter representing the coverage of the *X* circular code or the *X* motifs in genes. Unexpectedly, this parameter identifies several relations between the *X* circular code and translation efficiency and/or kinetics. We first investigate whether a correlation exists between the *X* circular code and the ‘optimal’ codons/dicodons associated with increased gene translation efficiency and mRNA stability. Then, we examine the recent evidence resulting from high-throughput technologies such as ribosome profiling, and demonstrate that the presence of *X* motifs in genes can be used as a predictor of gene expression level. Taken together, these observations provide evidence supporting the idea that motifs from the *X* circular code represent a new genetic signal, participating in the maintenance of the correct reading frame and the optimization and regulation of gene expression.

## Results

In this section, we first compare the 20 codons of the *X* circular code with the optimal codons and dicodons that have been shown to influence translation efficiency. Then, using previously published experimental data, we investigate whether a correlation exists between the presence of *X* motifs in genes and the level of gene expression.

### *X* codons correlate with optimal codons

We compared the 20 codons that belong to the *X* circular code with the ‘codon optimality code’ resulting from various statistical and experimental studies in metazoan [31], as well as in *S. cerevisiae* [32]. In these studies, the codon stabilization coefficient (CSC) was used as a robust and conserved measure of how individual codons contribute to shape mRNA stability and translation efficiency. Fig 3 shows the mean ranking of optimal codons, according to the CSC score, from four different experiments (in *S. cerevisiae*, zebrafish, *Xenopus* and *Drosophila*), where the highest ranking codon is the most optimal one. The *X* codons are ranked significantly higher than non-*X* codons (i.e. the 41 coding codons which do not belong to the circular code *X*), according to a Mann-Whitney signed rank test (*z*-score = 4.3, *p*-value < 0.00001). In other words, optimal codons for mRNA stability and elongation rate are significantly enriched in *X* codons.

**Fig 3.**
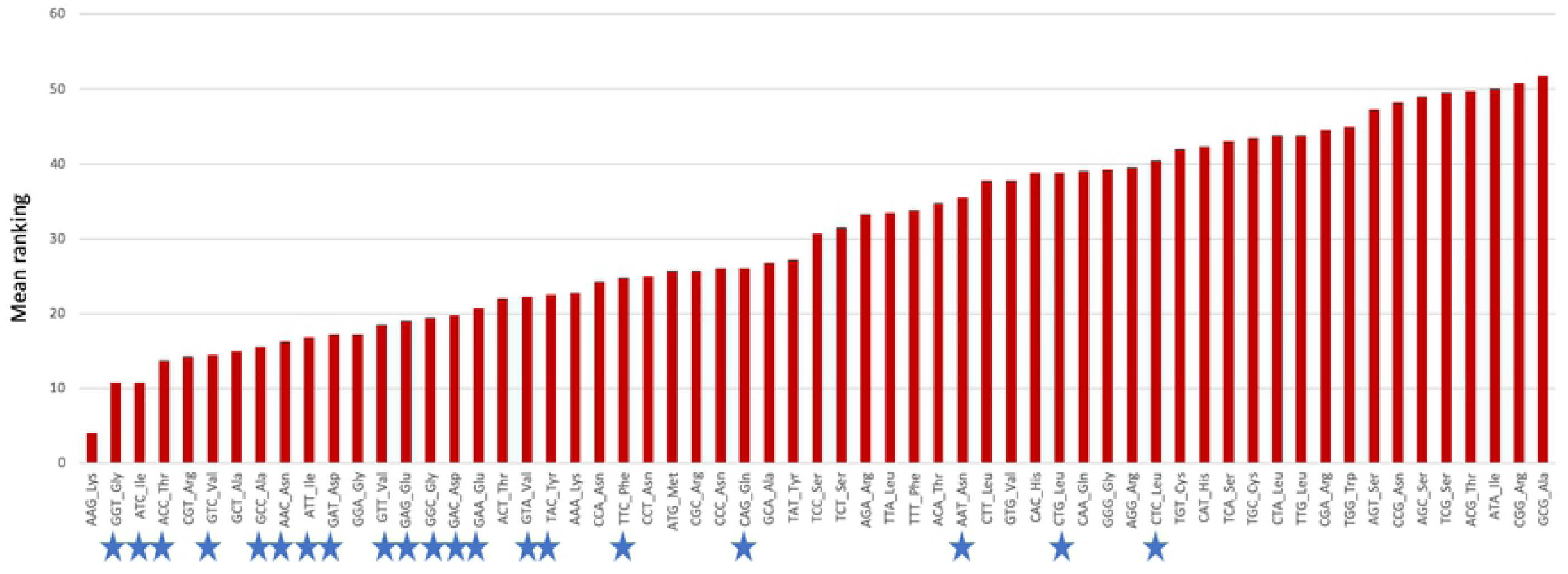
Optimal codons for translation elongation rate and mRNA stability in different eukaryotic species (*S. cerevisiae*, zebrafish, *Xenopus* and *Drosophila*). Codons are ordered according to their mean ranking obtained in four different experiments. Codons belonging to the *X* code are identified by a blue star.

### *X* codons correlate with the dicodons associated with increased expression

In recent years, emerging evidence has shown that translational rates may be encoded by dicodons rather than single codons [33-35]. For example, a large-scale screen in *S. cerevisiae* [33] assessed the degree to which codon context modulates eukaryotic translation elongation rates beyond effects seen at the individual codon level. The authors screened yeast cell populations housing libraries containing random sets of triplet codons within an ORF encoding superfolder Green Fluorescent Protein (GFP). They found that 17 dicodons were strongly associated with reduced GFP expression, i.e. associated with a substantial reduction of the translation elongation rate. This set included the known inhibitory dicodon *CGA*-*CGA* and was enriched for codons decoded by wobble interactions. Of these 17 dicodons associated with slower translation elongation rates, none are composed of 2 *X* codons (Table 1).

**Table 1.** Dicodons enriched in low or high abundance proteins, in two different studies: Gamble [33] and Diambra [34]. *X* codons are highlighted in blue.

| Dicodons associated with low abundance |  | Dicodons associated with high abundance |  |
| --- | --- | --- | --- |
| Gamble | Diambra | Diambra |  |
| AGG-CGA | AAA-ATA | **AAC-AAC** | **GAC-ACC** |
| AGG-CGG | **AAT**-GCA | **AAC-AAG** | **GAC-TAC** |
| ATA-CGA | **AAT**-TGG | **AAC-ACC** | **GAT-GCT** |
| ATA-CGG | AGT-AAG | AAG-TCC | **GCC-AAC** |
| CGA-ATA | AGT-GTG | **ACC-AAC** | **GCC-AAG** |
| CGA-CCG | ATA-**GGT** | **ACC-AAG** | **GCC-ACC** |
| CGA-CGA | **ATT**-AAA | **ACC-ACC** | **GCC-ATC** |
| CGA-CGG | CAA-AGT | **ACC-ATC** | **GCC-GCC** |
| CGA-**CTG** | **CAG**-AAA | **ACC-ATT** | **GGT-GTC** |
| CGA-GCG | **GAA**-AGT | **ACC-GCC** | **GTC-AAG** |
| **CTC**-CCG | **GAA**-CTA | **ACC-TTC** | **GTC-ACC** |
| **CTG**-ATA | GCA-TTT | **ATC-AAC** | **GTC-ATC** |
| **CTG**-CCG | TAT-AAA | **ATC-AAG** | **GTT-GCC** |
| **CTG**-CGA | TAT-CCG | **ATC-ACC** | **TAC-AAC** |
| **GTA**-CCG | TTT-**CAG** | **ATC-ATC** | **TAC-AAG** |
| **GTA**-CGA | TTT-TTT | **ATT-GCC** | **TCC-ACC** |
| GTG-CGA |  | CCA-CCA | **TTC-AAC** |
|  |  | CGT-CGT | **TTC-AAG** |
|  |  | **GAC-AAC** | **TTC-ACC** |
|  |  | **GAC-AAG** | **TTC-ATC** |

A subsequent statistical analysis of coding sequences of nine organisms [34] identified dicodons with significant different frequency usage for coding either lowly or highly abundant proteins. The working hypothesis was that sequences encoding abundant proteins should be optimized, in the sense of translation efficiency. 16 dicodons were identified with a preference for low abundance proteins, while 40 dicodons presented a preference for high abundance proteins. None of the 16 dicodons associated with low abundance proteins are composed of 2 *X* codons (Table 1). In contrast, 27 of the 40 dicodons associated with high abundance proteins correspond to 2 *X* codons, and only 3 dicodons do not contain any *X* codons (Table 1).

These recent studies support the idea that codons in coding sequences are likely arranged in an organized way, and that the local sequence context contributes to the effects of codon usage bias on gene regulation. Strikingly, our observations support the hypothesis that codon context may be linked in some way to the *X* circular code. In the next section, we describe more detailed analyses that test this hypothesis further.

### *X* motifs are enriched in the minimal gene set

Based on the increasing evidence of the importance of codon context [35-39], we hypothesized that if the *X* circular code plays a role in gene regulation, then we might expect to see a non-random use, or ‘clusters’, of *X* codons along the length of the gene. In previous work [23-24], we defined an *X* motif as a series of consecutive *X* codons (of length at least 4 codons in order to always retrieve the reading frame, see also Materials and Methods) in a gene sequence and searched for such *X* motifs in the reading frames of different genes. This approach allowed us to demonstrate that the reading frames of genes in yeasts and in mammals are significantly enriched in such *X* motifs. To test the hypothesis that the *X* motifs represent a more universal signature, we analyzed a set of 81 genes that were previously defined as a ‘minimal gene set’ [40]. At that time, the ‘minimal gene set’ genes were found to be conserved in all species. We used the *Mycoplasma genitalium* genes provided in the original study, as well as 15,822 orthologous sequences (5503 eukaryotes, 9205 bacteria and 1114 archaea), and identified all *X* motifs in the reading frame with a minimum length of 4 codons. Fig 4 shows the density *d*_*𝒮*_*(X)*(defined in Equation (1)) of the *X* motifs in the mRNA sequences. To evaluate the significance of the enrichment, as in previous work [23-24], we used a randomization model in which we generated N=100 random codes that preserved most of the properties to the *X* code, except the circularity. We then identified all random motifs from the 100 random codes and calculated mean values for the 100 codes.

**Fig 4.**
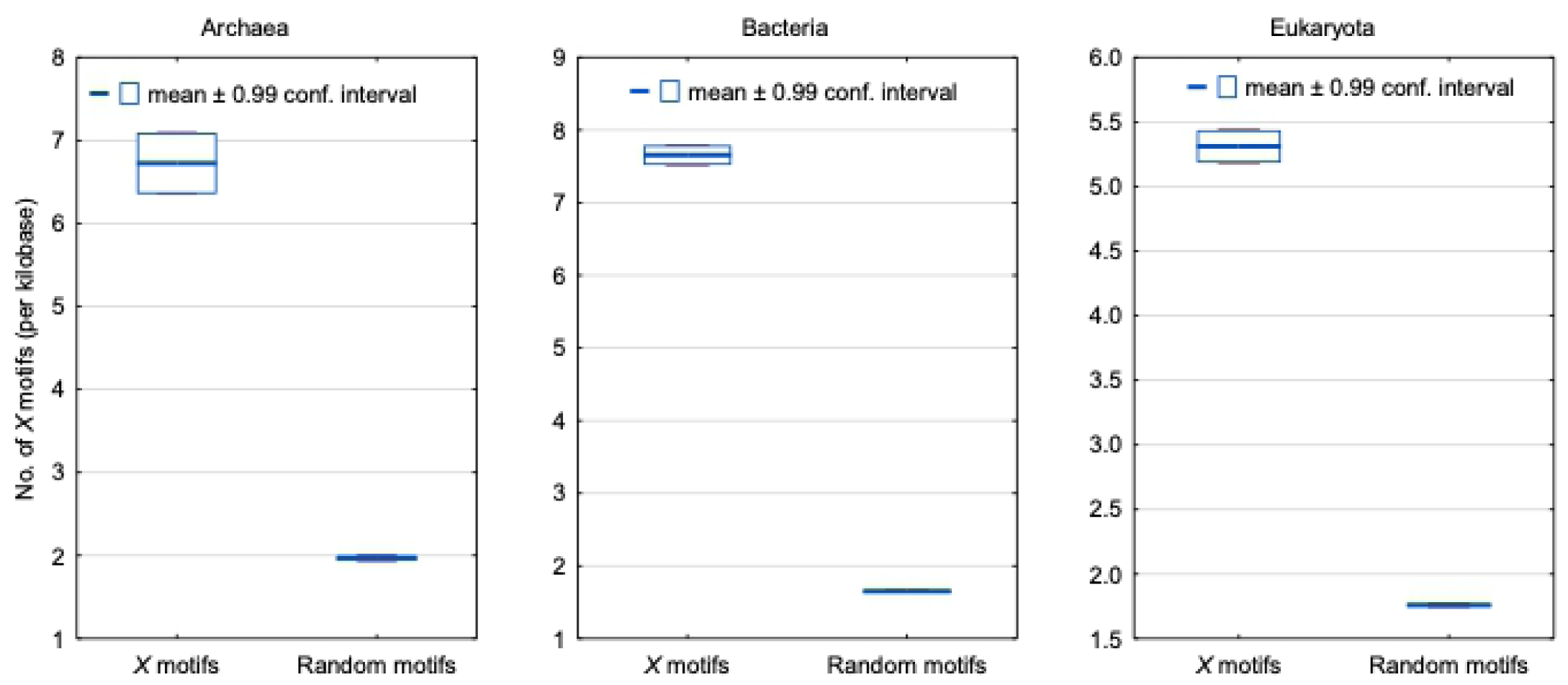
Number of *X* motifs (per kilobase; density *d*_*𝒮*_*(X)*defined in Equation (1)) in the mRNA sequences of the ‘minimal gene set’. The distributions of the number of *X* motifs identified in the sequences from the three domains of life are indicated by boxplots representing the mean number with a ±0.99 confidence interval. The distributions of the number of *R* random motifs (see Materials and Methods) identified in the same sequences are shown for statistical evaluation. There is a very strong statistical significance as confirmed by a one-sided Student’s *t*-test with a *p*-value *p* < 10^−100^ for each set of sequences from archaea, bacteria and eukaryota.

The density of *X* motifs found in the minimal gene set sequences belonging to the three domains of life, is significantly higher than the density of random motifs according to a one-sided Student’s *t*-test (*p* < 10^−100^) for each set of sequences from archaea, bacteria and eukaryota. This study demonstrates that *X* motifs are significantly enriched in the minimal gene set, and seem to be a universal feature of gene sequences in all three domains of life.

### *X* motifs are enriched in codon-optimized genes

If *X* motifs modify the codon usage in favor of optimal codons for translational efficiency, then we would expect that increasing the number of *X* motifs in a gene would increase the expression level. In an indirect way, we have shown that this is indeed the case. We previously showed that synthetic genes, which were re-designed for optimized protein expression, generally have more *X* motifs [24]. S1A Fig shows an example of the protein L1h from human papillomavirus (HPV-16), optimized for expression in mammalian cell lines and leading to significantly increased expression [41]. Here, the wild type gene contains only 3 *X* motifs, while the optimized gene construct has a total of 21 *X* motifs. It is important to note that classical codon optimization strategies do not always increase protein expression levels. S1B Fig shows another example involving the L1s protein from human papillomavirus (HPV-11) optimized for expression in the potato *Solanum tuberosum* [42]. In this case, only a low level of L1 expression was observed for the codon-optimized gene. In this example, we did not observe a significant difference between the number of *X* motifs in the wild type and optimized sequences (5 *X* motifs in the wild type gene compared to 4 in the optimized construct).

Codon replacement strategies have also been applied to the design of attenuated viruses, although in this case frequent codons are replaced with rare ones. Using quantitative proteomics and RNA sequencing, the molecular basis of attenuation in a strain of bacteriophage T7 (*Escherichia coli* K-12) was investigated [43]. The authors engineered the *E. coli* major capsid protein gene (gene 10A) to carry different proportions of suboptimal, rare codons. Transcriptional effects of the recoding were not observed, but proteomic observations revealed that translation was halved for the completely recoded major capsid gene, with subsequent effects on virus fitness (measured as doublings/hour). We obtained the sequences with 10%, 20%, 30% and 50% recoding from [44] and identified the density *d*_*𝒮*_*(X)*(defined in Equation (1)) of *X* motifs in each construct. Fig 5 clearly shows the correlation between the fitness obtained for each recoded sequence and the density of *X* motifs observed. The authors suggested that recoding of gene 10A reduced capsid protein abundance probably by ribosome stalling rather than ribosome fall-off.

**Fig 5.**
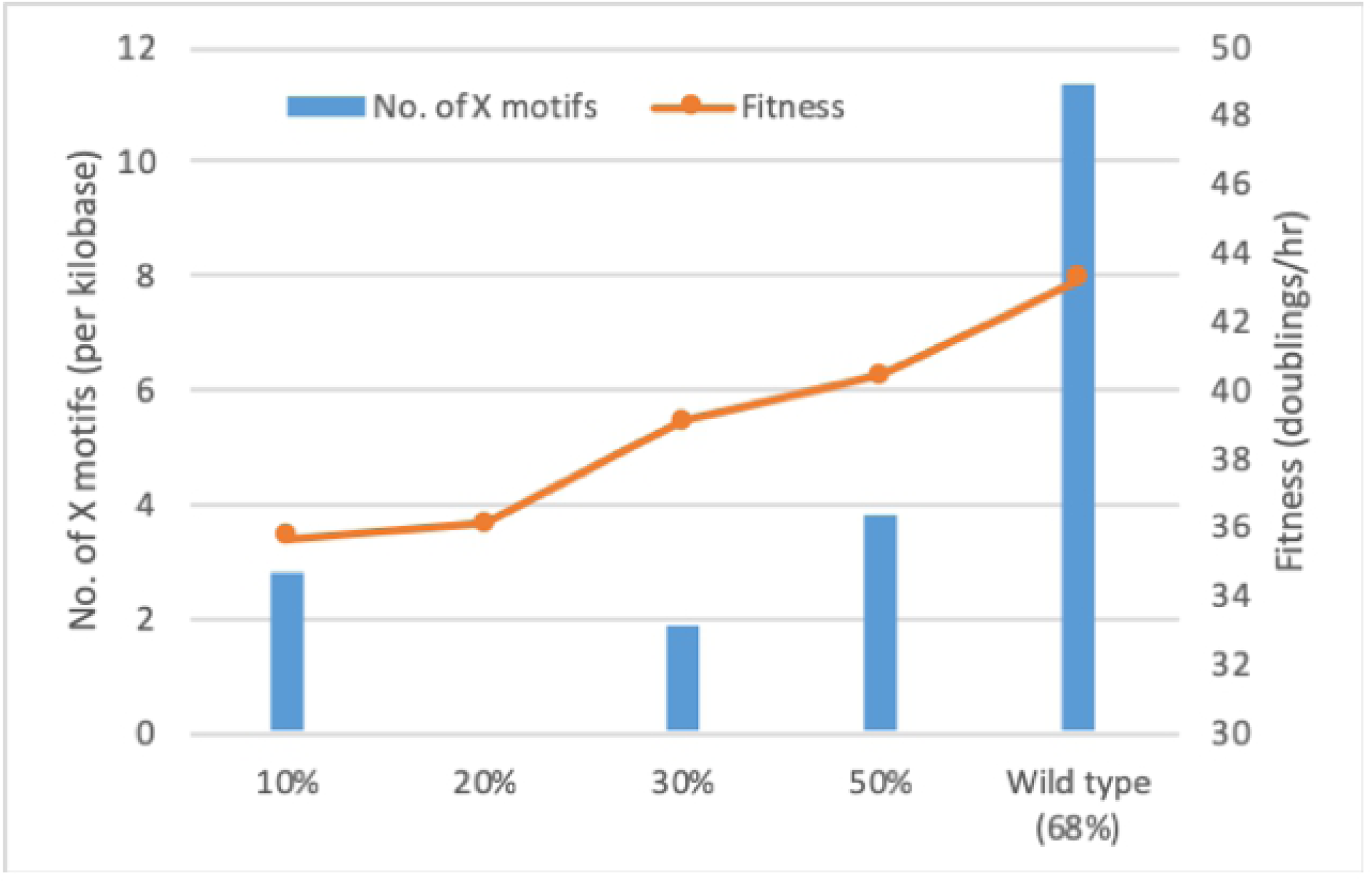
Histogram of the number of *X* motifs (per kilobase; density *d*_*𝒮*_*(X)*defined in Equation (1)) in the recoded version of the gene 10A from *Escherichia coli* K-12, compared to the wild type sequence. The orange plot indicates the viral fitness values corresponding to each construct.

In general, codon optimization is a successful strategy for improving protein expression in heterologous systems. However, simply replacing all rare codons by frequent codons can have negative effects *in vivo* [45]. Rare codons have the potential to slow down the translation elongation rate, due to the relatively long dwell time of the ribosome while searching for rare tRNAs. Several studies have suggested that gene-wide codon bias in favor of slowly translated codons serves as a regulatory means to obtain low expression levels of protein when desired, for example, in the case of regulatory genes, or where excess of the protein may be detrimental or lethal to the cell. An example, in *Neurospora crassa*, demonstrated that codon optimization of the central clock protein FRQ actually abolished circadian rhythms [46]. Different optimized constructs of the wild type gene *frq* were used in the study, where either the N-terminal end (codons 1-164) or the middle region (codons 185-530) was optimized. All optimized constructs gave higher levels of FRQ protein, this led to a different structural conformation. The density of *X* motifs (defined in Equation (1)) identified in the different wild type and optimized constructs is shown in Table 2. As in the previous examples, the optimized constructs contain significantly more *X* motifs (for instance, density of 10.2 in the N-terminal end of the fully optimized construct compared to 4.1 in the wild type). This example shows how non-optimal codon usage, and the associated reduction in the number of *X* motifs, can be used *in vivo* to regulate protein expression and to achieve optimal protein structure and function.

**Table 2.** Comparison of *X* motifs (per kilobase; density *d*_*𝒮*_*(X)*defined in Equation (1)) in the wild type gene *frq* and different optimized constructs for the *Neurospora crassa* FRQ protein. In the ‘mid opt’ constructs, only the non-preferred codons were changed; for ‘full opt’ constructs, every codon was optimized.

| Region | *frq* construct | Nb of $X$ motifs (per kilobase) |
| --- | --- | --- |
| N-terminal<br>(1-164) | wild type | 4.1 |
|  | mid opt | 6.1 |
|  | full opt | 10.2 |
| Middle<br>(185-530) | wild type | 3.9 |
|  | full opt | 5.8 |

In nature, the translation efficiency of a gene may vary at different conditions, cell types and tissues [47-50]. Thus, it has been proposed that the codon optimization should take into account other factors in addition to replacing rare codons by frequent ones, a process termed ‘codon harmonization’ [51-53]. Taken together, the examples described above suggest that it may be important for such harmonization strategies to consider the effect of codon replacement on the insertion or deletion of *X* motifs.

### *X* motifs correlate with translation efficiency and mRNA stability

We have shown previously that the reading frames of genes in *S. cerevisiae* are significantly enriched in *X* motifs [16]. Since then, ribosomal profiling has enabled a more detailed study of translation efficiency for a large set of 5450 genes from this organism [54]. A central assumption of ribosome profiling is that indirect measurement of the kinetics of translation *via* ribosome footprint occupancy on transcripts is directly reflective of true protein synthesis. The authors thus estimated the average translation rate of each gene, using experimental measurements of ribosome occupancy. Again, we identified the *X* motifs in the complete set of 5450 genes and calculated the density of *X* motifs (defined in Equation (1)) in three subsets of the genes having different estimated translation rates (Fig 6). We observed that genes with higher translation rates had significantly more *X* motifs than those with lower translation rates. The density of *X* motifs is higher for the sequences with medium translation rates than for those with low translation rates (one-sided Student’s t-test *p* < 10^−10^) and for the sequences with high translation rates than for those with medium translation rates (one-sided Student’s t-test *p* < 10^−14^). This result demonstrates the link between the total time needed for ribosome transition on a mRNA and density of *X* motifs along the length of the sequence.

**Fig 6.**
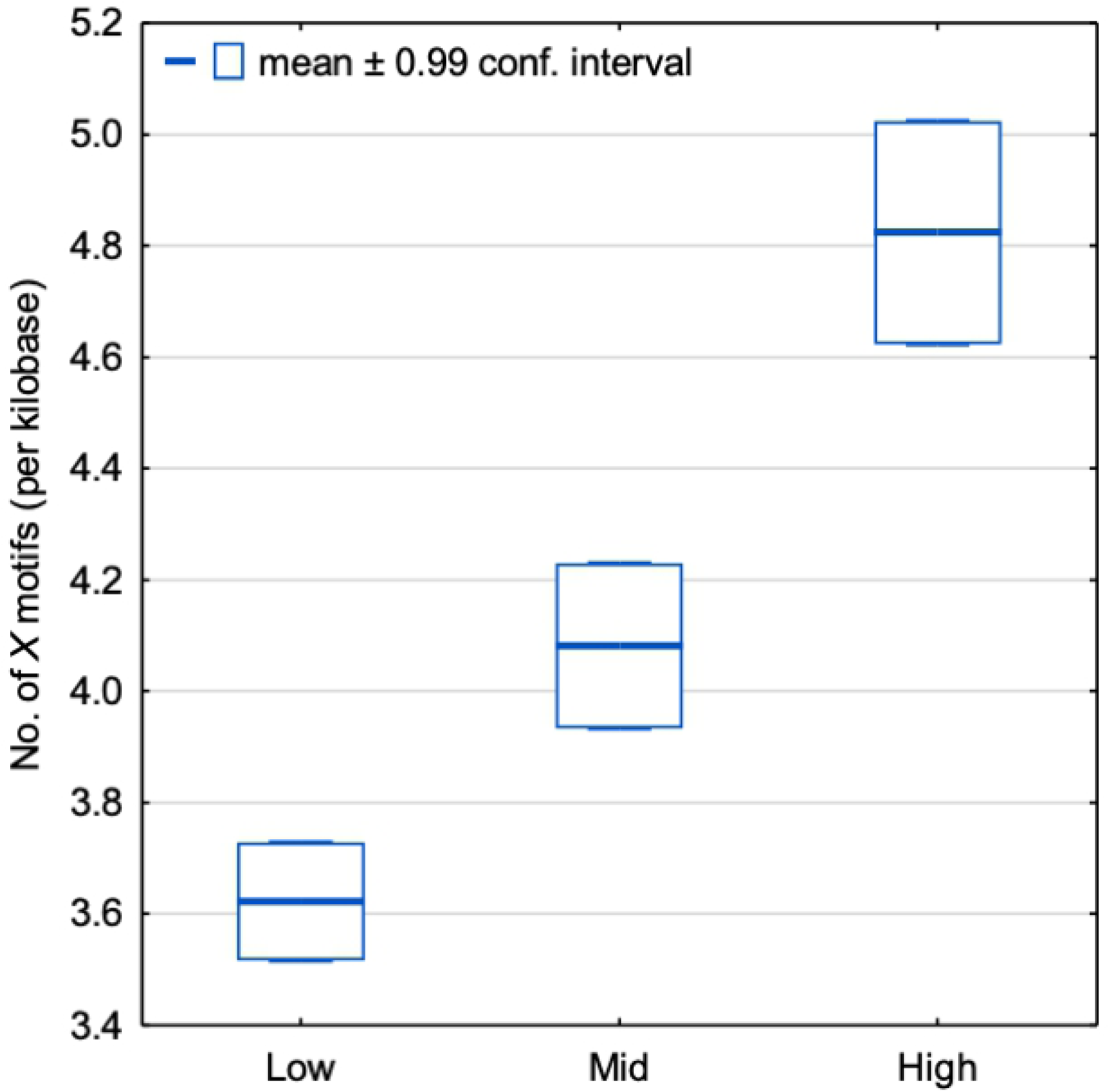
Number of *X* motifs (per kilobase; density *d*_*𝒮*_*(X)*defined in Equation (1)) for *S. cerevisiae* genes: 1323 genes with low translation rates (estimated translation rate < 0.03), 1378 genes with medium translation rates (estimated translation rate 0.05-0.09) and 1324 genes with high translation rates (estimated translation rate > 1.1). The distributions of the number of *X* motifs identified in the genes are indicated by boxplots representing the mean number with a ±0.99 confidence interval. The statistical significance is confirmed by two one-sided Student’s t-tests with: *p* < 10^−10^ between the sequences with medium translation rates and those with low translation rates; and *p*<10^−14^ between the sequences with high translation rates and those with medium translation rates.

To investigate whether *X* motifs might play a role in modulating ribosome speed in specific regions in mRNA, we considered single protein studies, where local translation elongation rate has been studied in detail. The first example concerns the study of a gene in *S. cerevisiae*, to investigate the link between translational elongation and mRNA decay [55]. In this study, various HIS3 protein constructs (length of 699 nucleotides) were designed with increasing codon optimality (measured by the CSC index) from 0% to 100%. We identified *X* motifs in the different constructs as before and compared them to the experimentally measured mRNA half-life. As the authors point out, the mRNA half-life is largely determined by the codon-dependent rate of translational elongation, since mRNAs whose translation elongation rate is slowed by inclusion of non-optimal codons are specifically degraded. The density of *X* motifs ranges from 0 in the 0% optimized construct to more than 7 in the 100% optimized sequence (Fig 7). The results suggest that the introduction of individual *X* motifs in specific regions can be used to increase the mRNA half-life.

**Fig 7.**
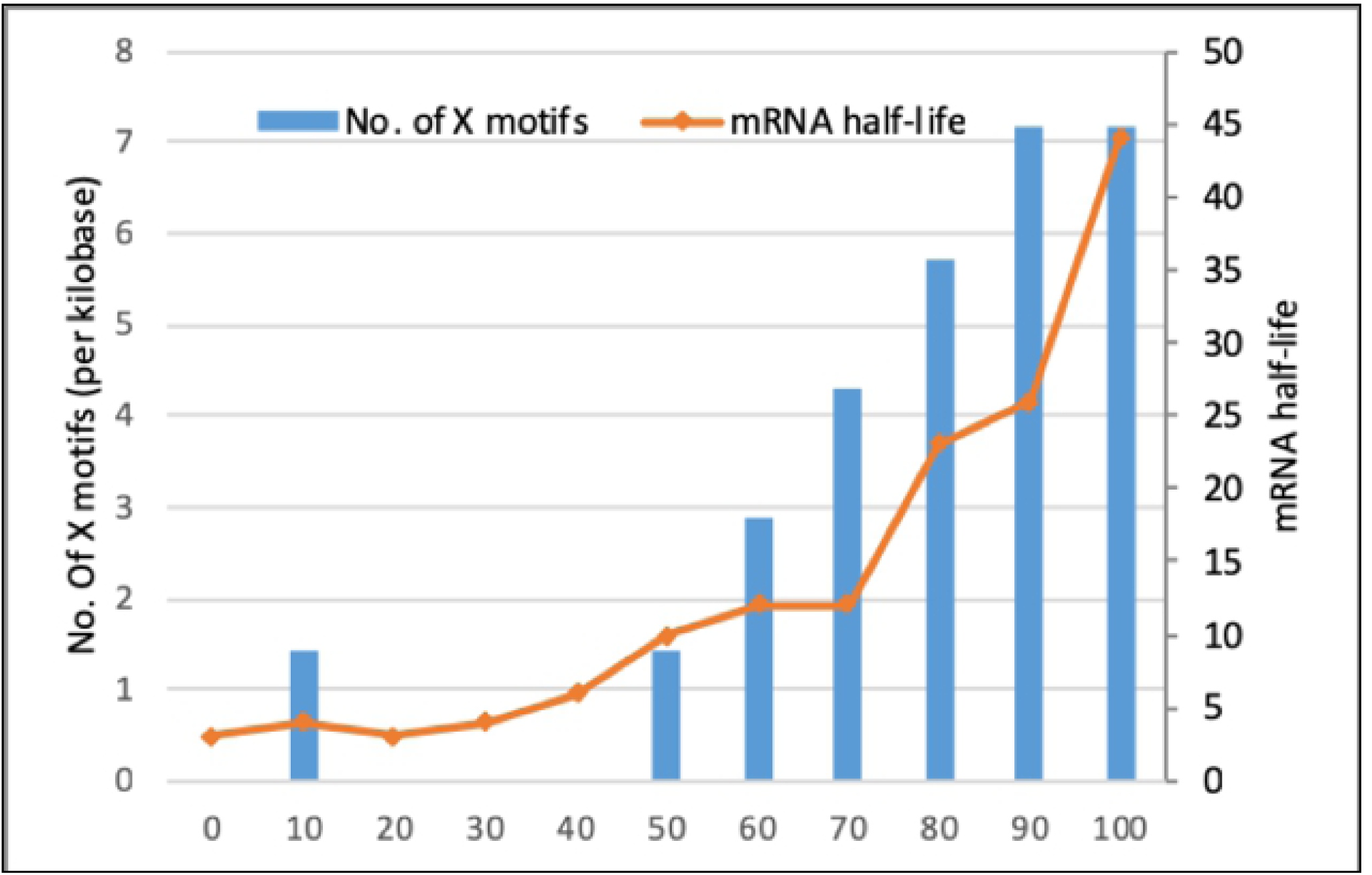
Histogram of the number of *X* motifs (per kilobase; density *d*_*𝒮*_*(X)*defined in Equation (1)) for different constructs corresponding to the *S. cerevisiae* HIS3 gene with 0-100% optimized codons. The orange plot indicates the mRNA half-life values corresponding to each construct.

The second example concerns a *Drosophila* cell-free translation system that was used to directly compare the rate of mRNA translation elongation for different luciferase constructs with synonymous substitutions [56]. The OPT construct was designed with the most preferred codons in all positions except for the first 10 codons, while the dOPT construct had the least preferred codons in all positions. The N-OPT, M-OPT and C-OPT constructs were created by replacing the N-terminal part (codons 11– 223), middle part (codons 224–423) and C-terminal part (codons 424–550) of the dOPT sequence with the corresponding optimized sequence, respectively. For each construct, the authors measured the time when the luminescence signal was first detected after start of translation. The time of first appearance (TFA) should thus reflect the speed of translation process. Higher TFA values were observed for each construct in the order dOPT < C-OPT < M-OPT < N-OPT < OPT, correlating well with an increasing density of *X* motifs (Fig 8). These results suggest that the introduction of *X* motifs in different regions of the gene significantly increased the rate of translation elongation, probably by speeding up ribosome movement on the mRNA.

**Fig 8.**
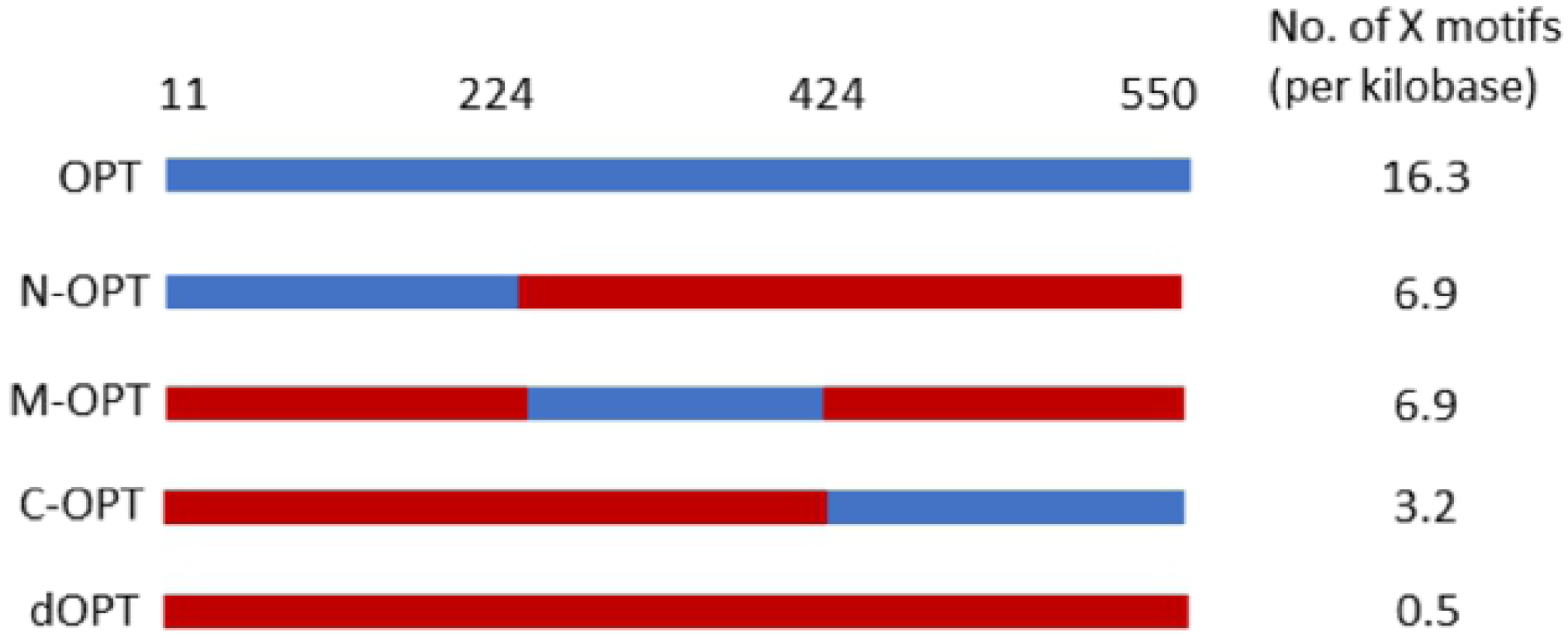
Number of *X* motifs (per kilobase; density *d*_*𝒮*_*(X)*defined in Equation (1)) in the different constructs corresponding to the *Drosophila* luciferase gene. Sequence regions shown in blue are codon optimized, and in red are the wild type sequence. The numbers above the sequences indicate the codon positions of the optimized regions.

We have highlighted the potential effects of *X* motifs on translation elongation speed, protein folding and function. The examples selected include studies in very different organisms, including viruses, fungi and insects with different codon usage bias (codon usage tables for these organisms are provided in S1 Table), but in all the examples a strong correlation is observed between ‘optimal’ codons and *X* codons. Taken together, the results support the idea that the use of *X* motifs is a conserved mechanism from viruses to animals that may participate in the modulation or regulation of the translation elongation rate along the mRNA.

## Discussion

In this work, we have combined two very distinct research domains: gene translation through the genetic code and the theory of circular codes which allows two processes simultaneously: reading frame retrieval and amino acid coding. Our hypothesis is that at least two codes operate in genes: the standard genetic code, experimentally proved to be functional, and the *X* circular code that has been shown to be statistically enriched in genes. For the first time here, we shed light on a number of biological experimental results by using the definition of a very simple parameter to analyze the density of *X* motifs in genes, i.e. motifs from the circular code *X*.

We would first like to make some comments about the mathematical structure of these two codes. The standard genetic code consists of 60 codons coding for 19 amino acids, the start codon *ATG* that codes for the amino acid *Met* and establishes the reading frame, and three non-coding stop codons {*TAA,TAG,TGA*}. The genetic code has a weak mathematical structure: a surjective coding map for the 60 codons and an incomplete self-complementary property for the 60 codons (e.g. the complementary codon of *TTA* coding the amino acid *Leu* is the stop codon *TAA*) implying a non self-complementary property for the four start/stop codons. The circular code *X* consists of 20 codons coding for 12 amino acids and has a strong mathematical structure: circularity for retrieving the reading frame, a surjective coding map, a complete self-complementary property for the 20 codons, a *C*^3^ property, etc. (reviewed in [12-13]).

We propose that the theory of circular codes can be used to shed light on many of the observed phenomena related to optimal codons/dicodons and the effects of codon optimization on different factors of gene expression, from transcriptional regulation to translation initiation, retrieval of the open reading frame, translation elongation velocities, and protein folding. We showed that optimal codons at the species and gene levels correlate well with the 20 codons that define the *X* circular code. Importantly, the optimal codons identified in diverse species [31] that increase translation elongation rates and mRNA stability are significantly enriched in *X* codons. We then studied a number of published experiments that used recent technologies to perform more detailed investigations of codon usage along the length of a gene, which suggest that codon context and local clusters of optimal or non-optimal codons may represent important regulatory signals for translation bursts and pauses [6,57]. In all these experiments, increased translation efficiency correlates with the number of *X* motifs present in the gene sequences. These observations raise the question: do *X* motifs somehow orchestrate elongation rate? Since it is known that translational elongation rate is intimately connected to mRNA stability, it is also tempting to suggest that *X* motifs are linked to the universal code of codon-mediated mRNA decay proposed by Chen and Coller [58].

The theory of the *X* circular code will have practical implications for improving the prediction of gene expression levels based on the gene sequence. Most of the current codon usage measures are dependent on the studied organism and the chosen expression system. In contrast, the presence of *X* motifs represents a universal signature that is significantly correlated with increased expression. Our previous work has already shown that *X* motifs can predict functional versus dubious genes in yeast [23] and can be used for rational gene design [24].

Translation of mRNA by the ribosome is a universal mechanism, and the most parsimonious explanation for the observed correlation between the presence of *X* motifs and increased translation elongation rates is that *X* motifs are somehow recognized by the ribosome. It is known that codon usage has effects on the major steps of translation elongation, including codon-anticodon decoding and peptide bond formation [57], as well as translocation which can be slowed down by mRNA secondary structure elements, such as pseudoknots and stem-loops [59]. Our hypothesis that *X* motifs in mRNA are recognized by the ribosome is further supported by recent ribosome profiling experiments in *Neurospora crassa*, which suggest that codon optimization increases the rate of ribosome movement on mRNA [60], and by the observation that translation elongation and mRNA stability are coupled through the ribosomal A-site [61]. Interestingly, our previous work has identified *X* motifs in the anticodon region of multiple tRNA genes, as well as in important functional regions of the ribosomal rRNA including the decoding centre [25-27].

How could motifs from the *X* circular code work? If the decoding unit at the ribosome is the anticodon then the comma-free codes would immediately return to the reading phase while the general circular codes would have a delay associated with reading at most four codons (exactly 13 nucleotides). If the decoding unit at the ribosome is the anticodon with adjacent nucleotides then the general circular codes could also immediately return to the reading phase. Does the self-complementary property of the *X* circular code contribute to coordination between *X* motifs in mRNA and *X* motifs in tRNA and/or rRNA? So far we have mainly discussed the effects of codon choices on the throughput of translation, but changes in the translation elongation process can clearly affect translation fidelity and accuracy, reviewed in [62]. For example, clustering of rare codons could deplete cognate tRNAs, increasing the probability of a near- or non-cognate tRNA occupying the decoding site, and this probability could be reflected in the frequency of miss-incorporation. In this case, it has been shown that the standard genetic code minimizes the impact of the mutations on the translated protein [56]. Clustering of identical rare codons also increases the probability of a frameshift during translation. Ribosome stalling at *Lys* codons triggers ribosome sliding on successive *AAA* codons. When ribosomes resume translation, they may shift in an incorrect reading frame. The ribosomes translating in the -1 or +1 frame usually quickly encounter out-of-frame stop codons that result in termination. Again, it has been suggested that the genetic code might be in some way optimized for frameshift mutations [63]. Given the inherent error correcting properties of circular codes, it is possible that the *X* circular code may play a role in the synchronization of the correct reading frame.

In the future, we hope to show that the simple parameter defined in this work to estimate the coverage of *X* motifs in genes is a useful factor that should be taken into account in codon optimization strategies or other experimental approaches involving gene expression. We also plan to investigate more complex parameters linked to X motifs, such as localized density patterns within specific regions of the genes.

## Materials and Methods

### Definition of the *X* motif density parameter

We define an *X* motif *m*_*s*_*(X)*as a series of at least 4 consecutive codons of the circular code *X* (defined in the Introduction) in the reading frame of a gene sequence *s*. For example, *m*_*s*_*(X)= CAGGACTACGTCGAC* is an *X* motif since *CAG, GAC, TAC* and *GTC* are codons of *X*. It is important to remember that any *X* motif with at least 4 consecutive *X* codons always allows the reading frame to be retrieved. Let *N(m*_*s*_*(X))*be the number of *X* motifs *m*_s_*(X)*in a gene sequence *s* of nucleotide length *l*_*s*_. Then the density *d*_*s*_*(X)* of *X* motifs in a gene sequence *s* is defined by the number of *X* motifs per kilobase in *s*, i.e.

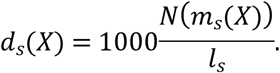

This density *d*_*s*_*(X)* in a sequence *s* can easily be extended to a density *d*_*𝒮*_*(X)*in a set *𝒮* of gene sequences *s* by dividing the total number of *X* motifs with the total nucleotide length, i.e.

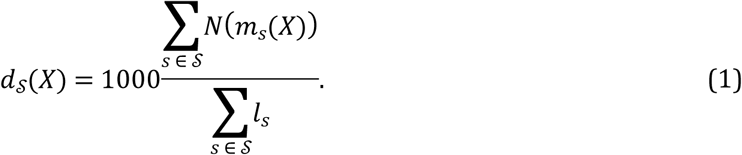

These densities *d*_*s*_*(X)* and *d*_*𝒮*_*(X)* are normalized parameters allowing to compare the coverage of *X* motifs in sequences of different lengths and in sequence populations of different sizes.

In order to evaluate the statistical significance of the obtained results, we also define 100 random codes *R* with similar properties to the *X* circular code, using the method described in Dila et al. (2019a). We then identified *R* random motifs *m*_*s*_(*R*) from these codes in the gene sequences and calculated the densities *d*_*s*_(*R*) and *d*_*𝒮*_(*R*) of *R* motifs, as for the *X* motifs.

### Data sources for analysis of optimal codons

As a measure of the optimality of each codon, we used the codon stabilization coefficient (CSC), defined by [31] as the Pearson correlation coefficient between the occurrence of each codon and the half-life of each mRNA. Thus, codons found more frequently in genes with longer mRNA half-lives have higher CSC values. The 61 coding codons can then be ranked according to their CSC scores in different organisms. We obtained the CSC rankings for each codon from previous studies in four species: zebrafish ([31], *Xenopus* [31], *Drosophila* [64] and *S. cerevisiae* [32]. We then calculated the mean CSC ranking for each codon in these four species.

Dicodons associated with reduced protein expression in *S. cerevisiae* were taken from a previous study [33]. Dicodons associated with low abundance or high abundance proteins were obtained from a previous study [34].

### Minimal gene set analysis

The minimal gene set of 81 genes conserved in all species was obtained from a previous study [40]. We used the *Mycoplasma genitalium* genes provided in the original study as a query, and searched for orthologues in the reference set of complete genomes for 317 species (144 eukaryotes, 142 bacteria and 31 archaea) in the OrthoInspector 3.0 database [65]. This resulted in a set of 15822 protein sequences (5503 eukaryotes, 9205 bacteria and 1114 archaea). For each protein sequence, we retrieved the mRNA sequences from the Uniprot database (www.uniprot.org) and identified all *X* motifs in the reading frame with a minimum length of 4 codons, using in-house developed software.

### Data sources for codon-optimized genes

Experimental data for synthetic genes re-designed for optimized protein expression were obtained from the SGDB database [66]. SGDB contains sequences and associated experimental information for synthetic (artificially engineered) genes from published peer-reviewed studies. We selected the gene entries where the synthetic sequence contained only synonymous codon changes, and where experimental protein expression levels were available for both the wild type and the synthetic gene. This resulted in a set of 45 gene pairs (wild type and synthetic gene), for which we identified all *X* motifs in the reading frame with a minimum length of 4 codons, using in-house developed software as before.

### Estimation of translation rates based on ribosome profiling data

Computational estimations of translation rates for 5450 *S. cerevisiae* genes were obtained from a previous study [54]. The authors performed a simulation of translation based on the totally asymmetric simple exclusion process (TASEP) model, using experimental measurements of the number of ribosomes on each transcript as well as RNA copy numbers to calibrate the parameters. For each of the 5450 gene sequences, we identified the *X* motifs using in-house developed software.

## Acknowledgements

This work was supported by Institute funds from the French Centre National de la Recherche Scientifique and the University of Strasbourg. The authors would like to thank the BISTRO and BICS Bioinformatics Platforms for their assistance. This work was supported by French Infrastructure Institut Français de Bioinformatique (IFB) ANR-11-INBS-0013, and the ANR Grants Elixir-Excelerate: GA-676559 and RAinRARE: ANR-18-RAR3-0006-02.

## Supporting information captions

S1 Fig. *X* motifs in the wild type and codon-optimized sequences from the SGDB database. **A**. L1h gene from human papillomavirus. **B**. L1s gene from human papillomavirus. The *X* motifs in the wild type sequence are shown in blue, and in the codon optimized sequences in red.

S1 Table. Codon usage tables for the species used in the different studies described in the main text. Data are from the HIVE-CUTS database at https://hive.biochemistry.gwu.edu/. *X* codons are highlighted in blue.

## References

1. Crick FH, Barnett L, Brenner S, Watts-Tobin RJ. General nature of the genetic code for proteins. Nature. 1961;192: 1227–1232.

2. Eslami-Mossallam B, Schram RD, Tompitak M, van Noort J, Schiessel H. Multiplexing genetic and nucleosome positioning codes: A computational approach. PLoS One. 2016;11: e0156905.

3. Prakash K, Fournier D. Evidence for the implication of the histone code in building the genome structure. Biosystems. 2018;164: 49–59.

4. Baralle M, Baralle FE. The splicing code. Biosystems. 2018;164: 39–48.

5. Cakiroglu SA, Zaugg JB, Luscombe NM. Backmasking in the yeast genome: encoding overlapping information for protein-coding and RNA degradation. Nucleic Acids Research. 2016;44: 8065–8072.

6. Yu CH, Dang Y, Zhou Z, Wu C, Zhao F, Sachs MS, Liu Y. Codon usage influences the local rate of translation elongation to regulate co-translational protein folding. Mol Cell. 2015;59: 744–754.

7. Weatheritt RJ, Babu MM. Evolution. The hidden codes that shape protein evolution. Science. 2013;342: 1325–1326.

8. Maraia RJ, Iben JR. Different types of secondary information in the genetic code. RNA. 2014;20: 977–984.

9. Bergman S, Tuller T. Widespread non-modular overlapping codes in the coding regions. Physical Biology. 2020;1088: 1478-3975/ab7083.

10. Babbitt GA, Coppola EE, Mortensen JS, Ekeren PX, Viola C, Goldblatt D, Hudson AO. Triplet-based codon organization optimizes the impact of synonymous mutation on nucleic acid molecular dynamics. Journal Molecular Evolution. 2018;86: 91–102.

11. Arquès DG, Michel CJ. A complementary circular code in the protein coding genes. Journal of Theoretical Biology. 1996;182: 45–58.

12. Michel CJ. A 2006 review of circular codes in genes. Computer and Mathematics with Applications 2008;55: 984–988.

13. Fimmel E, Strüngmann L. Mathematical fundamentals for the noise immunity of the genetic code. Biosystems 2018;164: 86–198.

14. Crick FH, Griffith JS, Orgel LE. Codes without commas. Proceedings of the National Academy of Sciences USA. 1957;43: 416–421.

15. Nirenberg MW, Matthaei JH. The dependence of cell-free protein synthesis in E. coli upon naturally occurring or synthetic polyribonucleotides. Proceedings of the National Academy of Sciences USA. 1961;47: 1588–1602.

16. Michel CJ. The maximal *C*3 self-complementary trinucleotide circular code *X* in genes of bacteria, archaea, eukaryotes, plasmids and viruses. Life. 2017;7: 20.

17. Frey G, Michel CJ. Circular codes in archaeal genomes. Journal of Theoretical Biology 2003;223: 413–431.

18. Frey G, Michel CJ. Identification of circular codes in bacterial genomes and their use in a factorization method for retrieving the reading frames of genes. Computational Biology and Chemistry 2006;30: 87–101.

19. Grosjean H, Westhof E. An integrated, structure- and energy-based view of the genetic code. Nucleic Acids Research. 2016;44: 8020–8040.

20. Michel CJ, Thompson JD. Identification of a circular code periodicity in the bacterial ribosome: origin of codon periodicity in genes? RNA Biology. 2020;17: 571–583.

21. Fimmel E, Michel CJ, Pirot F, Sereni J-S, Strüngmann L. Mixed circular codes. Mathematical Biosciences 2019;317: 108231.

22. El Soufi K, Michel CJ. Circular code motifs in genomes of eukaryotes. Journal of Theoretical Biology. 2016;408: 198–212.

23. Michel CJ, Ngoune VN, Poch O, Ripp R, Thompson JD. Enrichment of circular code motifs in the genes of the yeast *Saccharomyces cerevisiae*. Life. 2017;7.

24. Dila G, Michel CJ, Poch O, Ripp R, Thompson JD. Evolutionary conservation and functional implications of circular code motifs in eukaryotic genomes. Biosystems. 2019a;175: 57–74.

25. Michel CJ. Circular code motifs in transfer RNAs. Computational Biology and Chemistry. 2013;45: 17–29.

26. Michel CJ. Circular code motifs in transfer and 16S ribosomal RNAs: a possible translation code in genes. Computational Biology and Chemistry. 2012;37: 24–37.

27. Dila G, Ripp R, Mayer C, Poch O, Michel CJ, Thompson JD. Circular code motifs in the ribosome: a missing link in the evolution of translation? RNA 2019b;25: 1714–1730.

28. Demongeot J, Seligmann H. Spontaneous evolution of circular codes in theoretical minimal RNA rings. Gene 2019a;705: 95–102.

29. Demongeot J, Moreira A. A possible circular RNA at the origin of life. Journal of Theoretical Biology. 2007;249: 314–324.

30. Demongeot J, Seligmann H. The uroboros theory of life’s origin: 22-nucleotide theoretical minimal RNA rings reflect evolution of genetic code and tRNA-rRNA translation machineries. Acta Biotheoretica. 2019b;67: 273–297.

31. Bazzini AA, Del Viso F, Moreno-Mateos MA, Johnstone TG, Vejnar CE, Qin Y, Yao J, Khokha MK, Giraldez AJ. Codon identity regulates mRNA stability and translation efficiency during the maternal-to-zygotic transition. EMBO Journal. 2016;35: 2087–2103.

32. Hanson G, Coller J. Codon optimality, bias and usage in translation and mRNA decay. Nature Reviews Molecular Cell Biology 2018;19: 20–30.

33. Gamble CE, Brule CE, Dean KM, Fields S, Grayhack EJ. Adjacent codons act in concert to modulate translation efficiency in yeast. Cell. 2016;166: 679–690.

34. Diambra LA. Differential bicodon usage in lowly and highly abundant proteins. PeerJ. 2017;5: e3081.

35. Guo FB, Ye YN, Zhao HL, Lin D, Wei W. Universal pattern and diverse strengths of successive synonymous codon bias in three domains of life, particularly among prokaryotic genomes. DNA Research. 2012;19: 477–485.

36. Clarke TF, Clark PL. Rare codons cluster. PLoS One. 2008;3: e3412.

37. Brar GA. Beyond the triplet code: Context cues transform translation. Cell. 2016;167: 1681–1692.

38. Chevance FFV, Hughes KT. Case for the genetic code as a triplet of triplets. Proceedings of the National Academy of Sciences USA. 2017;114: 4745–4750.

39. Sharma AK, Sormanni P, Ahmed N, Ciryam P, Friedrich UA, Kramer G, O’Brien EP. A chemical kinetic basis for measuring translation initiation and elongation rates from ribosome profiling data. PLOS Computational Biology. 2019;15: e1007070.

40. Koonin EV. How many genes can make a cell: the minimal-gene-set concept. Annual Review of Genomics and Human Genetics. 2000;1: 99–116.

41. Leder C, Kleinschmidt JA, Wiethe C, Müller M. Enhancement of capsid gene expression: Preparing the human papillomavirus type 16 major structural gene L1 for DNA vaccination purposes. Journal of Virology. 2001;75: 9201–9209.

42. Warzecha H, Mason HS, Lane C, Tryggvesson A, Rybicki E, Williamson AL, Clements JD, Rose RC. Oral immunogenicity of human papillomavirus-like particles expressed in potato. Journal of Virology. 2003;77: 8702–8711.

43. Jack BR, Boutz DR, Paff ML, Smith BL, Bull JJ, Wilke CO. Reduced protein expression in a virus attenuated by codon deoptimization. G3 (Bethesda). 2017;7: 2957–2968.

44. Bull JJ, Molineux IJ, Wilke CO. Slow fitness recovery in a codon-modified viral genome. Molecular Biology Evolution. 2012;29:2997–3004.

45. Gingold H, Pilpel Y. Determinants of translation efficiency and accuracy. Molecular Systems Biology. 2011;7: 481.

46. Zhou M, Guo J, Cha J, Chae M, Chen S, Barral JM, Sachs MS, Liu Y. Non-optimal codon usage affects expression, structure and function of FRQ clock protein Nature. 2013;495: 111–115.

47. Charneski CA, Hurst LD. Positively charged residues are the major determinants of ribosomal velocity. PLoS Biology. 2013;11: e1001508.

48. Gardin J, Yeasmin R, Yurovsky A, Cai Y, Skiena S, Futcher B. Measurement of average decoding rates of the 61 sense codons in vivo. Elife. 2014;3.

49. Weinberg DE, Shah P, Eichhorn SW, Hussmann JA, Plotkin JB, Bartel DP. Improved ribosome-footprint and mRNA measurements provide insights into dynamics and regulation of yeast translation. Cell Rep. 2016;14: 1787–1799.

50. Wu CC, Zinshteyn B, Wehner KA, Green R. High-resolution ribosome profiling defines discrete ribosome elongation states and translational regulation during cellular stress. Molecular Cell. 2019;73: 959-970.e5.

51. Brule CE, Grayhack EJ. Synonymous codons: Choose wisely for expression. Trends in Genetics. 2017;33: 283–297.

52. Villada JC, Brustolini OJB, Batista da Silveira W. Integrated analysis of individual codon contribution to protein biosynthesis reveals a new approach to improving the basis of rational gene design. DNA Research. 2017;24: 419–434.

53. Mignon C, Mariano N, Stadthagen G, Lugari A, Lagoutte P, Donnat S, Chenavas S, Perot C, Sodoyer R, Werle B. Codon harmonization - going beyond the speed limit for protein expression. FEBS Letters. 2018;592: 1554–1564.

54. Diament A, Feldman A, Schochet E, Kupiec M, Arava Y, Tuller T. The extent of ribosome queuing in budding yeast. PLOS Computational Biology. 2018;14: e1005951.

55. Boël G, Letso R, Neely H, Price WN, Wong KH, Su M, Luff J, Valecha M, Everett JK, Acton TB, Xiao R, Montelione GT, Aalberts DP, Hunt JF. Codon influence on protein expression in E. coli correlates with mRNA levels. Nature. 2016;529: 358–363.

56. Blazej P, Wnetrzak M, Mackiewicz D, Mackiewicz P. Optimization of the standard genetic code according to three codon positions using an evolutionary algorithm. PloS one. 2018;13: e0201715.

57. Rodnina MV. The ribosome in action: Tuning of translational efficiency and protein folding. Protein Science. 2016;25: 1390–1406.

58. Chen YH, Coller J. A universal code for mRNA stability? Trends in Genetics. 2016;32:687–688.

59. Wu B, Zhang H, Sun R, Peng S, Cooperman BS, Goldman YE, Chen C. Translocation kinetics and structural dynamics of ribosomes are modulated by the conformational plasticity of downstream pseudoknots. Nucleic Acids Research. 2018;46: 9736–9748.

60. Zhou Z, Dang Y, Zhou M, Li L, Yu CH, Fu J, Chen S, and Liu Y. Codon usage is an important determinant of gene expression levels largely through its effects on transcription Proceedings of the National Academy of Sciences USA. 2016;113: E6117–E6125.

61. Hanson G, Alhusaini N, Morris N, Sweet T, Coller J. Translation elongation and mRNA stability are coupled through the ribosomal A-site. RNA. 2018;24: 1377–1389.

62. Liu Y, Sharp J.S, Do DH, Kahn RA, Schwalbe H, Buhr F, Prestegard JH. Mistakes in translation: Reflections on mechanism. PLoS One. 2017;12: e0180566.

63. Geyer R, Madany Mamlouk A. On the efficiency of the genetic code after frameshift mutations. PeerJ. 2018;6: e4825.

64. Burow D.A, Martin S, Quail J.F, Alhusaini N, Coller J, Cleary MD. Attenuated codon optimality contributes to neural-specific mRNA decay in drosophila. Cell Reports. 2018;24: 1704–1712.

65. Nevers Y, Kress A, Defosset A, Ripp R, Linard B, Thompson JD, Poch O, Lecompte O. OrthoInspector 3.0: open portal for comparative genomics. Nucleic Acids Research. 2019;47: D411–D418.

66. Wu G, Zheng Y, Qureshi I, Zin H.T, Beck T, Bulka B, Freeland SJ. SGDB: a database of synthetic genes re-designed for optimizing protein over-expression. Nucleic Acids Research. 2007;35: D76–D79.

